# Behavioral gain following isolation of attention

**DOI:** 10.1101/2020.12.17.423241

**Authors:** Grace Edwards, Anna Berestova, Lorella Battelli

## Abstract

Stable sensory perception is achieved through balanced excitatory-inhibitory interactions of lateralized sensory processing. In real world experience, sensory processing is rarely equal across lateralized processing regions, resulting in continuous rebalancing. Using lateralized attention as a case study, we predicted rebalancing lateralized processing following prolonged spatial attention imbalance could cause a gain in attention the opposite direction. In neurotypical human adults, we isolated covert attention to one visual field with a 30-minute attention-demanding task and found an *increase* in attention in the opposite visual field after manipulation. We suggest a gain in lateralized attention in the previously unattended visual field is due to an overshoot through attention rebalancing. The offline post-manipulation effect is suggestive of long-term potentiation affecting behavior. Our finding of visual field specific attention rebound could be critical for the development of clinical rehabilitation for patients with a unilateral lesion and lateralized attention deficits. This proof-of-concept study initiates the examination of overshoot following the release of imbalance in other lateralized control and sensory domains, important in our basic understanding of lateralized processing.

## Introduction

Avoiding hazards while driving, catching a ball, and spotting your friend in a busy restaurant, involve careful deployment of lateralized visual, attentional, motor and tactile processing. Left and right hemispheric regions control right- and left-sided processing of exogenous input and endogenous control, respectively. Here, we examine the outcome of prolonged imbalance in lateralized processing.

Short-term imbalance between right and left lateralized processing causes an inhibitory interaction between lateralized regions, prioritizing processing in the activated hemisphere. For example, left limb tactile stimulation inhibits somatosensory processing of the right limb tactile stimulation (Palmer et al., 2012) and inhibitory non-invasive brain stimulation to left attention regions increases right hemispheric attention processing (Hilgetag et al., 2001). Studies on patients with unilateral lesions provide a window into severe and ongoing lateralized processing imbalance. Deficits in left visual field attention are typically found in patients with a right frontal-parietal unilateral lesion. Kinsbourne (1977) suggested that attentional neglect of the contralesional visual field result from a combination of the lesion and a hyperactivation of the healthy hemisphere, which further inhibits the lesioned cortex. The hyperactivity of the healthy cortex is hypothesized to be in response to a lack of inhibition from the lesioned cortex (Corbetta et al., 2005; Kinsbourne, 1977). However, patient studies are unable to detail how lateralized processing is rebalanced *after* a period of imbalance, which occurs often in the healthy cortex. For example, when driving on the outside lane of a highway, we fixate on the road ahead, but attend to the right visual field continuously for hazards. How does this prolonged imbalance effect subsequent attention processing across the visual field?

Studies on monocular deprivation (i.e. prolonged covering of one eye) have provided some insight to the product of prolonged imbalanced in sensory processing, albeit within lateralized sensory processing regions, rather than between. Lunghi et al., (2011) found prolonged monocular deprivation increased the processing strength of visual input to the deprived eye, post-deprivation. Examination of GABA concentration in a follow-up study indicated a reduction of inhibition between the ocular dominance columns of the left and right eye in the early visual cortex (Lunghi et al., 2015). Therefore, with long-lasting sensory imbalance, the commonly found inhibitory interaction between the eyes was muted, resulting in a rebound processing of sensory input to the deprived eye.

Here we determine if rebound can occur between lateralized processing regions using the case study of lateralized visual attention. Forty-two participants first performed bilateral Multiple Object Tracking (MOT) to obtain a baseline attention performance in the left and right visual fields (Figure 1a). MOT measures covert attention as participants track target objects in amongst distractor objects in the left and right visual fields separately, whilst maintaining central fixation. Next, participants were randomly assigned to one of three experimental procedures (Figure 1b): Group 1) performed right unilateral MOT whilst maintaining fixation and ignoring MOT stimuli on the left. This isolated their attention to the right visual field for 30 minutes. Group 2) performed left unilateral MOT while ignoring the MOT stimuli on the right, isolating their attention left for 30 minutes. Control Group 3) performed 30 minutes of bilateral MOT, tracking targets on both left and right. Immediately after the manipulation all participants performed bilateral MOT. Following prolonged attention isolation, we expected a rebound in tracking performance in the ignored visual field for Group 1 and 2.

**Figure 1:**
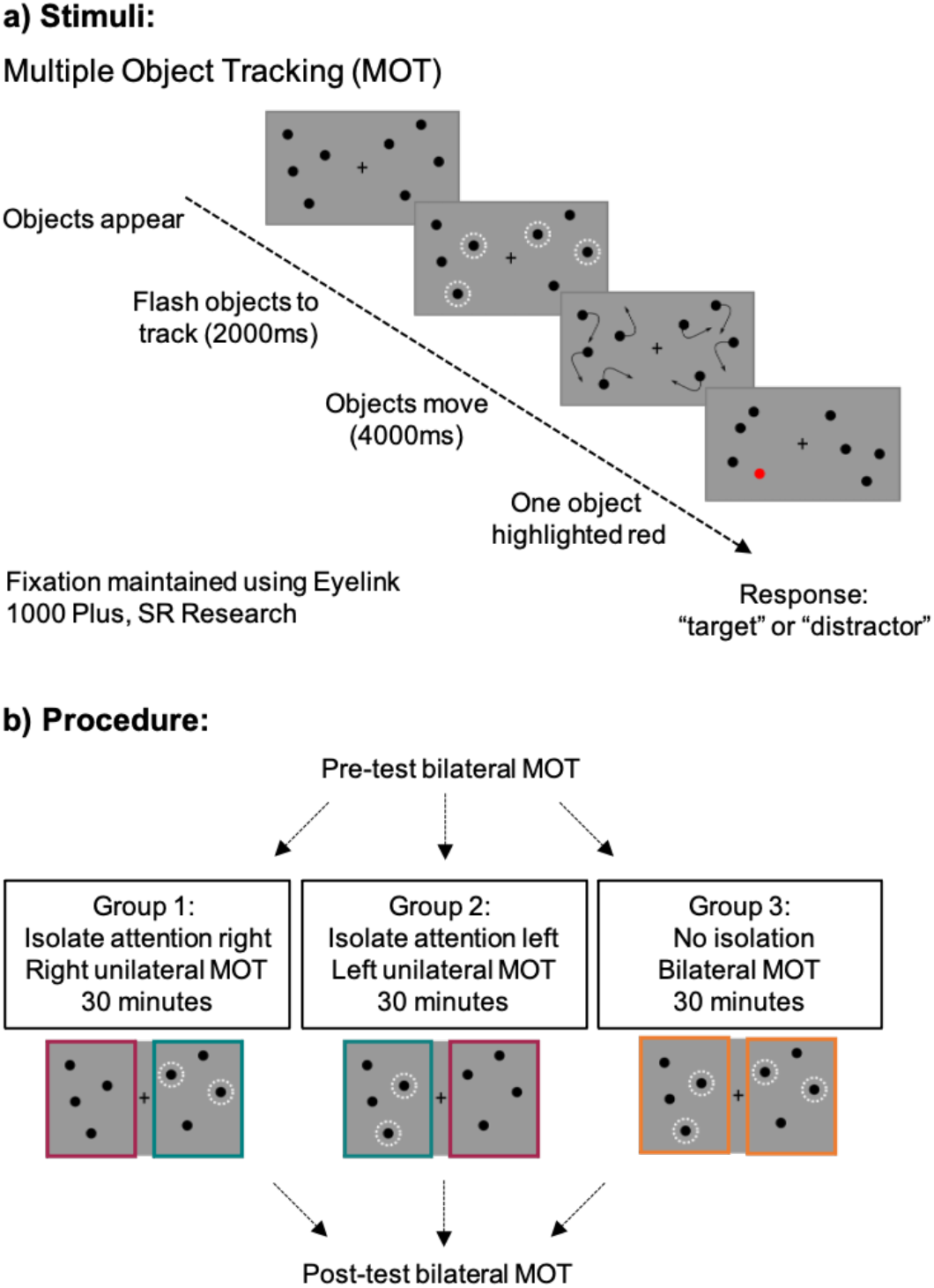
**a) Bilateral Multiple Object Tracking (MOT) stimuli.** Visual field specific attention measured using bilateral MOT or unilateral MOT (depicted in 1b). Fixation maintained using gaze-contingent programming with Eyelink 1000, SR Research. **b) Procedure.** Unilateral MOT employed to isolate attention to one visual field for 30 minutes in Groups 1 and 2. The difference from pre- to post-test bilateral MOT is determined for the Attended Visual Field in teal, Ignored Visual Field in maroon, and in the control visual fields in orange.

## Results

### Isolation manipulation impact on tracking performance

We first examined the impact of prolonged tracking in one visual field on modulation of visual field specific attention from pre- to post-isolation. We found a main effect of Session (χ^2^(1)=7.8866, p=0.0050, *glmer*) and no main effect of Visual Field (χ^2^(1)=3.2210, p=0.0727, *glmer*) or Manipulation (χ^2^(2)=1.0618, p=0.5881, *glmer*). We also found no three-way interaction between Session, Visual Field and Manipulation (χ^2^(2)=1.9270, p=0.3816, *glmer*). However, in examining the two-way interactions, we found an interaction between Manipulation and Session (χ^2^(2)=9.4680, p=0.0088, *glmer*; Figure 2a). This interaction indicates a significant change in tracking from pre- to post-manipulation as a function of attention isolation, however this interaction was not different across left and right visual field (Figure 2b). Model comparisons support Manipulation x Session interaction as the best and least complex fit for our data (see Supporting Information).

**Figure 2.**
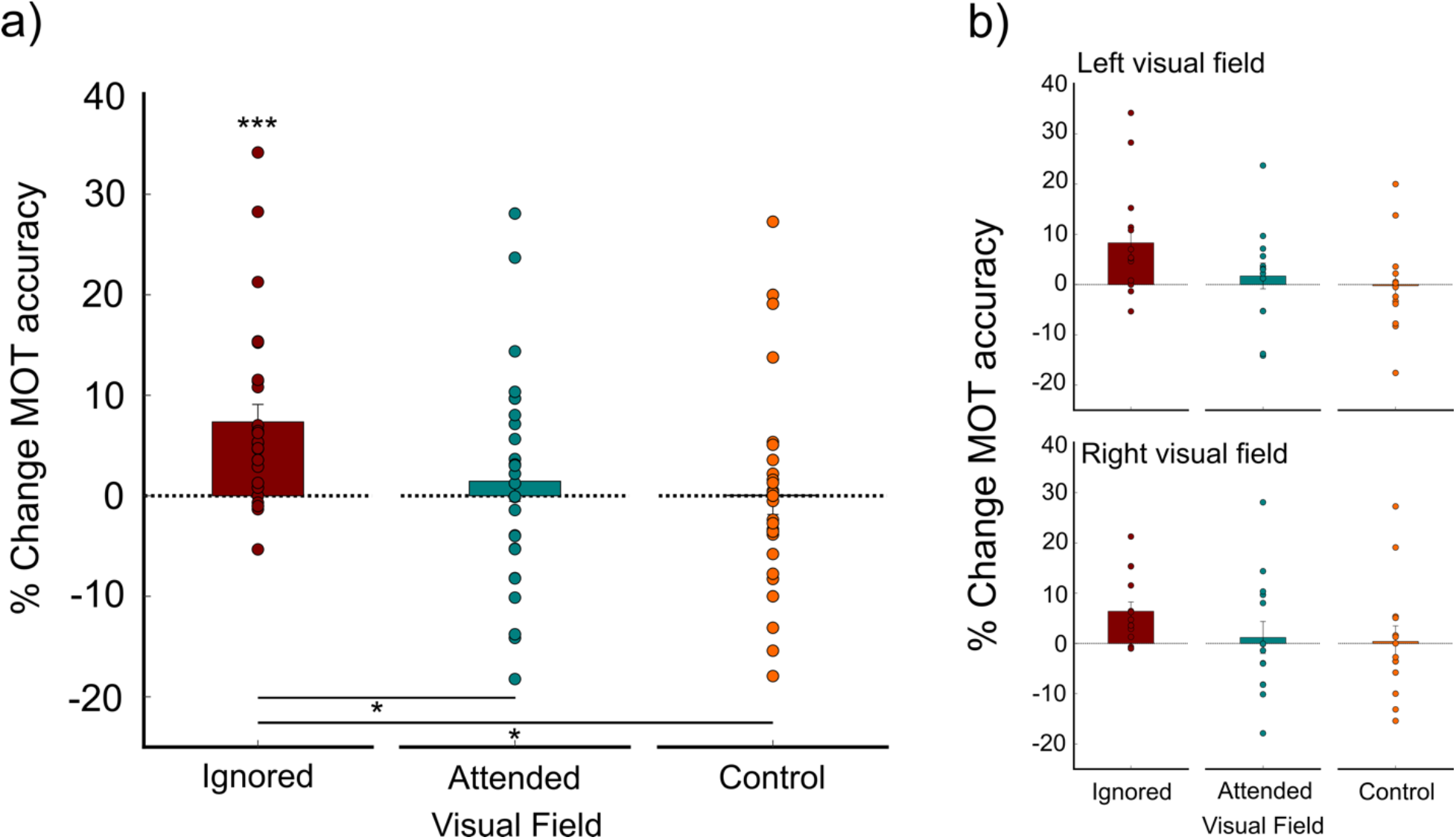
Change in MOT accuracy due to intervention. a) Data collapsed across left and right visual fields. Ignored visual field in maroon, attended visual field in teal, control visual field with bilateral tracking in orange. *** above maroon bar indicates p=0.0001 difference from pre- to post-manipulation. * between Ignored and Control indicates a difference significant to p=0.0205. * between Ignored and Attended indicates a difference significant to p=0.0439. b) Data separated by visual field, demonstrating no difference if manipulation is performed in right or left visual field.

Collapsing across visual field, we found a significant increase in tracking performance from pre- to post-manipulation for the Ignored Visual Field (Figure 2a; Maroon bar, estimate=0.6683, se=0.164, z=4.079, p=0.0001, confidence interval (CI)=0.278, 1.059; *emmeans()* with *adjust “mvt”*). No change in tracking performance was found for the Attended Visual Field (Teal bar, estimate=0.2052, se=0.161, z=1.277, p=0.487, CI=−0.173,0.588; *emmeans()* with *adjust “mvt*) or for the Control (Orange bar, estimate=0.0244, se=0.190, z=0.128, p=0.9989, CI=-0.334,1.299; *emmeans()* with *adjust “mvt*).

Furthermore, we found a significant difference in tracking between the Ignored Visual Field and the Control (estimate=0.0727, se=0.0276, t.ratio=2.634, p=0.0205, CI=0.01,0.135; *emmeans(),* with *adjust “mvt”*) and between the Ignored and Tracked Visual Fields (estimate=0.0590, se=0.0251, t.ratio=2.353, p=0.0439, CI=0.001,0.117; *emmeans(),* with *adjust “mvt”*).

### Pre-manipulation and during-manipulation tracking performance

Importantly, we found no difference between tracked, ignored, and control visual fields prior to manipulation (p>0.05; *emmeans()* with *adjust “mvt”*; Figure S2; all individual p-values reported in the Supplemental Table). Furthermore, no difference was found between groups during the attention manipulation period. We split the data into 10-minute time-bins to examine how isolation impacted behavior across the 30-minute period. We found no main effect of group (χ^2^(2)=4.1535, p=0.1253, *glmer*), nor main effect of time-bin (χ^2^(2)=2.5838, p=0.2747, *glmer*), and no interaction between group and time-bin (χ^2^(4)=8.9881, p=0.0614, *glmer;* Figures S3 & S4).

## Discussion

Forcing attention to one visual field for a prolonged duration resulted in an increase in attention to the opposite visual field immediately after manipulation. The boost of attention suggests a rebound in behavior following prolonged imbalance between lateralized attention processing regions. Rebalancing of neural activity is controlled via homeostatic plasticity which compensates for changes in activity levels to maintain stability of neuronal excitability (Wu et al., 2020). The inactivated attention processing regions may benefit from homeostatic gain control in the attempt to equalize visual field attention post-manipulation.

Another plausible account of our findings is post-inhibitory rebound spiking, where neuronal rebound has been demonstrated following prolonged inhibition (Kuffler & Eyzaguirre, 1955). Prolonged lateralized attention recruits lateralized attention processing regions (Culham et al., 1998), which in turn inhibit cortical homologues processing the opposite visual field (Hilgetag et al., 2001). It is plausible our behavioral rebound is a result of post-inhibitory spiking which initiated long term potentiation (Pugh & Raman, 2006). Considering we record the impact of manipulation offline (up to ten minutes after), Hebbian plasticity fits our behavioral change well.

The post-inhibitory rebound interpretation of our data is in opposition to previous research examining intracortical sensory imbalance, which found reduced inhibition. In adults, imbalance of visual input through covering of one eye (monocular deprivation) causes an increase processing for the *deprived* eye (Lunghi et al., 2011). The strengthening of the deprived eye has parallels to our study, where we find the strengthening of attention to the unattended (or attention-deprived) visual field. Using magnetic resonance spectroscopy (MRS), Lunghi et al. (2015) demonstrated the strengthening of the deprived eye correlated with decreased GABAergic inhibition in the early visual cortex. The lack of inhibition described by Lunghi et al., 2015 indicated that post-inhibitory rebound may not have caused the attention gain found here. However, rebound activity with decreased inhibition has also been demonstrated when examining center-surround receptive fields in cat primary visual cortex (Ozeki et al., 2009), where rebound activity was mediated by lack of excitation of the center rather than increased inhibition from the surround.

Interestingly, six 2-hour sessions of monocular deprivation to the unhealthy eye can restore visual acuity in amblyopic adults lasting at least one year (Lunghi et al., 2019). The restoration of visual acuity was interpreted as a plastic change following multiple prolonged periods of decreased inhibition between the two eyes during monocular deprivation. Using right attention isolation, we may exacerbate the visual field attention imbalance in right lateralized stroke patients, which could cause rebound in the impaired left visual field. Perhaps with multiple sessions of right attention isolation, we could also cause a long-lasting plastic change improving left visual field attention in right lateralized stroke patients.

Our results demonstrated that thirty minutes of tracking in one visual field did not induce training within the attended field, and therefore could not account for a training transfer to the unattended visual field. Previous research has demonstrated multiple sessions of MOT are necessary for successful training (Strong & Alvarez, 2017).

Behavioral gain in the ignored visual field following attention isolation ignites many questions. Firstly, what is the underlying mechanism which supports the rebound? Follow-up MRS studies examining GABA and glutamate complex concentration following attention isolation would characterize the role of inhibition in our behavioral finding (Frangou et al., 2019). Secondly, is rebound following isolation a shared characteristic across other lateralized sensory and control networks? A ubiquitous rebound characteristic could be useful for promoting interventions for plastic change. Thirdly, following along the lines of intervention, could isolation be a useful tool for the rehabilitation of patients with unilateral stroke exhibiting lateralized deficits? Exacerbating the existing imbalance in sensory and control processing may cause post-isolation rebound favoring contralesional performance. The intervention would focus on the *healthy* hemisphere the driving recovery, with a clear benefit in the ease of the approach for stroke survivors. Finally, sensory specific attention has been demonstrated to suppress cortical processing of other senses (Frank et al., 2021; Iurilli et al., 2012), suggesting activity gain for the processing of other senses may be possible using prolonged attention isolation.

## Method

### Participants

Forty-six right-handed individuals residing in the Cambridge & Boston area of Massachusetts volunteered to take part in the experiment (26 females; age range 20-40 years). The study was approved by Harvard University’s Institutional Review Board: The Human Research Protection Program. All participants gave written informed consent. Four participants were excluded from analysis due to data recording errors, leaving 42 participants in total. All participants had normal or corrected-to-normal vision.

### Stimuli

Participants viewed the stimuli on a 24-inch LCD Dell screen at a distance of 50 cm (screen resolution: 1980 x 1200), run via a 2010 Apple Mac mini. All stimuli were presented using Psychtoolbox in Matlab (Brainard, 1997).

### Bilateral Multiple Object Tracking Paradigm

Bilateral multiple object tracking (MOT) is a well-established task for the recruitment of top-down attention (Pylyshyn & Storm, 1988). When participants perform bilateral MOT (in both left and right visual fields simultaneously), lateralized attention processing regions are activated in the right and left hemispheres (Culham et al., 1998). In bilateral MOT, each trial began with a fixation point presented centrally for 1000 ms on a gray background (luminance: 19.5 cd/m^2^). Participants were required to maintain central fixation for the duration of the experiment controlled via an Eyelink 1000 Plus. If participants moved their eyes outside of a 1.5° boundary box, the trial was immediately restarted (see *Eye-tracking Acquisition)*. Four objects (black discs, 1.89 cd/m^2^. radius 0.25°) were then presented either side of fixation (eight discs total). Two discs on either side of fixation flashed (at 2 Hz for 2 sec) to indicate the targets participants had to track for the duration of the trial. All discs (targets and distractors) then moved along random trajectories within a 6°-by-6° area, at a constant speed for 4000 ms. The speed of the objects was set according to individual subjects’ threshold (see *Thresholding* section). The closest an object could come to fixation on the left or right visual field was 2°. Each object repelled each other to maintain a minimum of 1.5° space, bouncing off the invisible boundaries, and never crossing the vertical midline. Once the objects stopped moving, one object was highlighted in red on either the left or right of fixation and the participant had to respond if the highlighted object was a “target” or “distractor” with a button press. Importantly, the left and right visual field were tested equally, randomly interleaved in the run. The participant was not aware of which visual field would be tested until the end of the trial, necessitating attention to targets in both visual fields throughout the duration of the task. After the button press, the fixation point turned green to indicate a correct answer, or red to indicate an incorrect answer. Each trial lasted 9.5 seconds total.

### Unilateral MOT during manipulation

Participants in the manipulation groups performed unilateral MOT during the manipulation period. Whilst fixating centrally and attending to stimuli in one visual field, lateralized attention processing regions are active in the contralateral hemisphere (Sheremata & Silver, 2015). Group 1) performed right unilateral MOT to isolate attention right and Group 2) performed left unilateral MOT to isolate attention left (Figure 1b). In unilateral MOT the participants were asked to track two objects in amongst two distractors in only one visual field for the duration of the manipulation while maintaining fixation within 1.5° of fixation (see *Eye-tracking Acquisition*). Four objects were also presented in the untested visual field to equalize visual field sensory input.

### Thresholding MOT speed

At the beginning of the experimental session, each participant performed the bilateral MOT task at different speeds to find the speed at which they performed at 75% correct (Figure 1b). At 75% correct participants perform below ceiling prior to manipulation, enabling examination of behavioral change due to the experimental manipulation. All subjects began with 16 trials of MOT to practice at the lowest possible speed (2°/sec, the easiest condition, see *Multiple Object Tracking Paradigm (MOT)* section for details). Following practice, seven test blocks were performed using a constant speed approach to thresholding. In each trial of the test block, the speed of the objects was randomly assigned between 2°-16° per second, with 16 trials per block. Therefore, the duration of thresholding was 20 minutes. Linear interpolation was used to determine the speed at which each individual performed at 75% correct (average threshold speed = 6.71 deg/sec; SD 2.68; Figure S1).

### Procedure

Following recruitment, each participant was randomly assigned to one of three groups. Across the three groups each participant performed the same experimental procedure, except during the manipulation phase (see Figure 1b). First, participants performed the MOT thresholding task to measure the speed at which each participant performed 75% correct (*see Thresholding MOT speed*). Then participants were required to perform 10 minutes of bilateral MOT at their fixed individual speed threshold to obtain a pre-manipulation performance baseline measure. Next, participants entered the manipulation phase. Participants in Group 1 isolated attention to the right visual field for 30 minutes performing unilateral MOT whilst maintaining central fixation (see *Unilateral Multiple Object Tracking Paradigm & Eye-tracking Acquisition*), while participants in Group 2 isolated attention to the left visual field for 30 minutes in an otherwise identical procedure to Group 1. Finally, participants in Group 3 performed 30 minutes of bilateral MOT, testing tracking performance in both the left and right visual field equally, thereby experiencing no isolation of attention to one visual field. Immediately after manipulation all participants performed 10 minutes of post-test bilateral MOT to determine if the manipulation had impacted MOT accuracy. In Group 1 and 2 we were interested in the modulation of attention in the ignored visual field after prolonged attention isolation (Highlighted in maroon in Figure 1). Group 3 served as a control for any learning effects following prolonged MOT task. On average, subjects performed 61 trials (standard deviation (SD): 6) in the pre-manipulation phase, 167 trials (SD: 28) during manipulation, and 60 trials (SD: 9) post-manipulation. Trial number varied due to participants ability to fixate centrally (see *Eye-tracking Acquisition*).

### Eye-tracking Acquisition

An Eyelink 1000 Plus eye-tracking system (SR Research) was used to ensure participants maintained fixation during each trial of the experiment. Calibration was performed at the beginning of each run. During the experiment, an invisible boundary box of 1.5° x 1.5° was placed around the central fixation point, any time a participant moved their eyes outside of the boundary box, the trial restarted. Participants could blink comfortably without restarting the trial.

### Behavioral Data Analysis

We first fit a general linear mixed effects model on participants multiple object tracking accuracy in R (R Core Team, 2019) using *glmer()* function from the *lme4* package (Bates et al., 2014). The between-subject predictor was Manipulation (Attended Visual Field, Ignored Visual Field, or Control) and the within-subject predictors were Session (Pre- or Post-Manipulation) and Visual Field (Left or Right). Following model comparisons, the formula for the selected model was: *glmer (Accuracy ~ Manipulation * Session * VF + (1+Session|Subs) + (1+VF|Subs), data = AttIso, family = binomial)*. The linear mixed effects model was trial based and random intercepts and slopes were included for each participant. We performed model comparisons to determine which interactions best fit the data. *Chi-squared* and *p-values* were reported for the interactions and main effects. Individual contrasts were performed using *emmeans()* and *adjust=”mvt”* for multiple comparisons (Lenth et al., 2020). Analysis script and data are available for download and review at: https://osf.io/3kus7/?view_only=4322dca7c5a64276a561acccfc9e3400

## Acknowledgements

Data collection funded by the Harvard Mind Brain Behavior Interfaculty Initiative. We thank John Assad, Patrick Cavanagh, Roger Strong and Hrag Pailian for helpful insights on a previous version of this manuscript.

## Contributions

GE & LB designed the experiment. GE & AB performed the experiments. GE analyzed the data. GE & LB wrote the manuscript.

