## Supplemental Information for "Behavioral gain following isolation of attention"

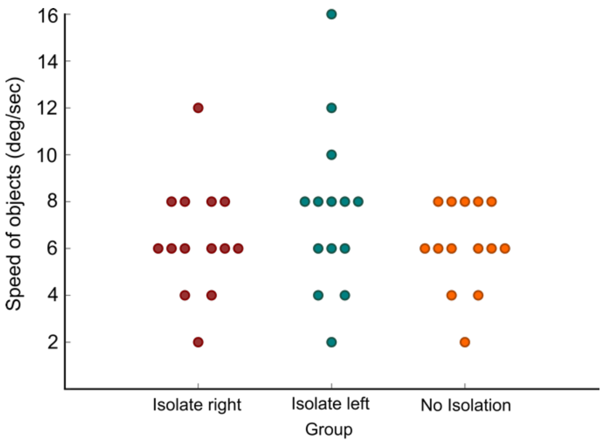

Supplemental Figure 1: Thresholding value per participant per group.

One-way ANOVA determined no difference in thresholding speed between the three groups (F(2,39)1.12, p=0.337).

### Model Comparisons

Model comparisons were conducted on various *glmer* models to determine the best model fit for the data collected.

### Syntax:

Mod1 <- glmer (Accuracy ~ Manipulation + Session + VF + *(1+Session|Subs) + (1+VF|Subs),* data = AttIso, family = binomial)
Mod2 <- glmer (Accuracy ~ Manipulation * Session + VF + *(1+Session|Subs) + (1+VF|Subs),* data = AttIso, family = binomial)
Mod3 <- glmer (Accuracy ~ Manipulation + Session * VF + *(1+Session|Subs) + (1+VF|Subs),* data = AttIso, family = binomial)
Mod4 <- glmer (Accuracy ~ Session + Manipulation * VF + *(1+Session|Subs) + (1+VF|Subs),* data = AttIso, family = binomial)
Mod5 <- glmer (Accuracy ~ Manipulation * Session * VF + *(1+Session|Subs) + (1+VF|Subs),* data = AttIso, family = binomial)

anova(Mod1, Mod2, Mod3, Mod4, Mod5)

| Mod | Df | AIC | BIC | loglik | deviance | Chisq | Chi Df | Pr(>Chisq) |
|---|---|---|---|---|---|---|---|---|
| Mod1 | 11 | 3540.1 | 3611.1 | -1759.1 | 3518.1 | | | |
| Mod3 | 12 | 3541.9 | 3619.3 | -1759.0 | 3517.9 | 0.2183 | 1 | 0.64350 |
| Mod2 | 13 | 3534.7 | 3618.5 | -1754.3 | 3508.7 | 9.2373 | 1 | 0.002371 |
| Mod4 | 13 | 3539.8 | 3623.7 | -1756.9 | 3513.8 | 0 | 0 | 1.00000 |
| Mod5 | 18 | 3538.5 | 3654.6 | -1751.3 | 3502.5 | 11.33005 | 5 | 0.045738 |

Model comparisons demonstrated Mod2 is the least complex and best fit for our data.

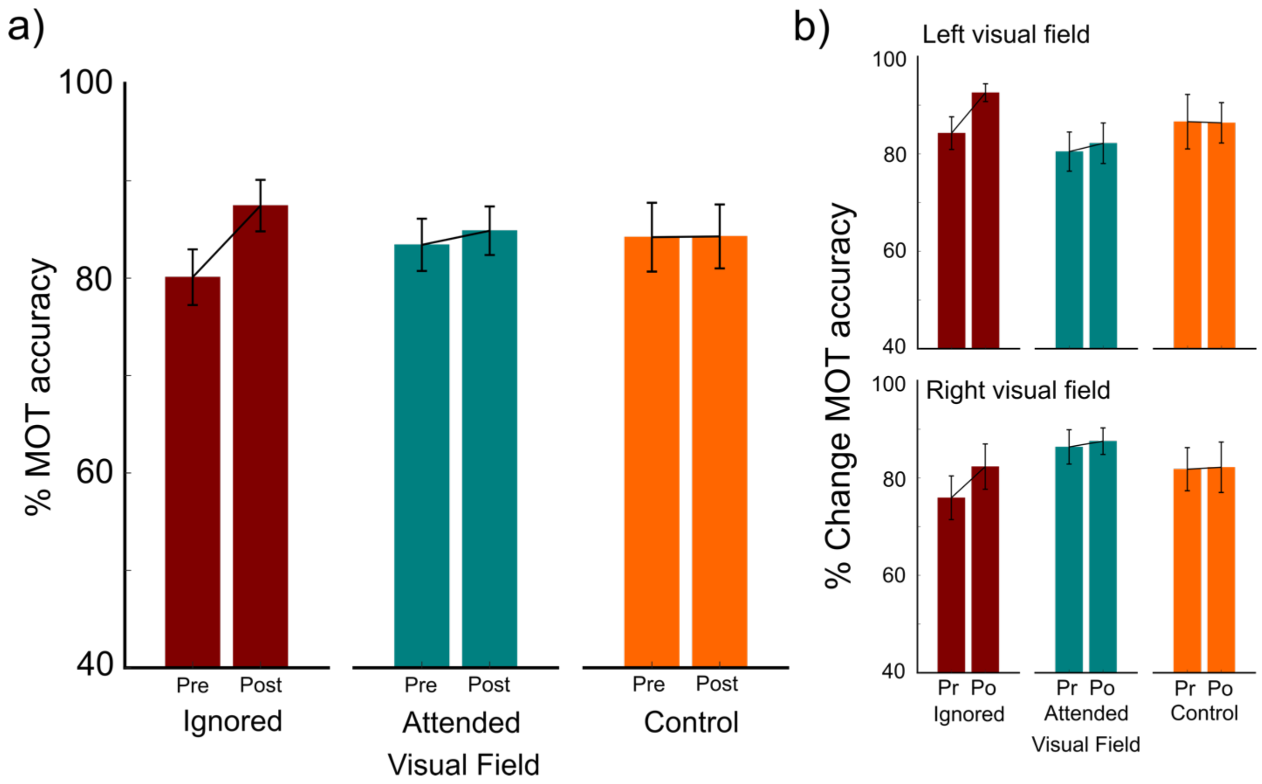


**Supplemental Figure 2: Percentage correct for pre- and post-manipulation**. a) Data collapsed across left and right visual fields. Ignored visual field in maroon, attended visual field in teal, control visual field with bilateral tracking in orange. Data separated by left and right visual field.

No difference found between group prior to intervention (p>0.05; *emmeans()* with *adjust "mvt"*).

| **Contrast** | **Estimate** | **SE** | **z.ratio** | **p.value** |
|---|---|---|---|---|
| Control > Ignored | 0.482 | 0.356 | 1.356 | 0.3451 |
| Ignored > Attended | -0.233 | 0.137 | -1.694 | 0.1922 |
| Attended > Control | -0.250 | 0.357 | -0.700 | 0.7512 |
| Control LVF > Ignored LVF | 0.605 | 0.438 | 1.381 | 0.3505 |
| Control LVF > Attended LVF | 0.841 | 0.436 | 1.929 | 0.1303 |
| Ignored LVF > Attended LVF | 0.236 | 0.410 | 0.575 | 0.8336 |
| Control RVF > Ignored RVF | 0.450 | 0.418 | 1.077 | 0.5283 |
| Control RVF > Attended RVF | -0.244 | 0.423 | -1.576 | 0.8329 |
| Ignored RVF > Attended RVF | -0.694 | 0.414 | -1.677 | 0.2138 |

**Supplemental Table: Comparisons between groups pre-manipulation.** Individual statistics for each comparison.

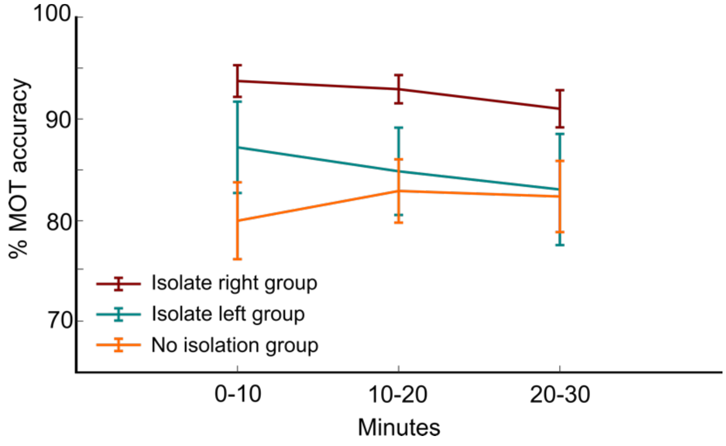


**Supplemental Figure 3: Percentage correct multiple object tracking accuracy during isolation period**. Data presented by group and split into 10-minute bins: Group 1 isolate attention right (maroon), therefore only right unilateral tracking, Group 2 isolate attention left (teal), therefore only left unilateral tracking, Group 3 no attention isolation (orange), therefore bilateral tracking.

We also determined if the decrease in tracking ability from the beginning to end of training could be used as an indication of deterioration in the trained visual field. Below we plot this metric against the impact of the intervention in the trained visual field.

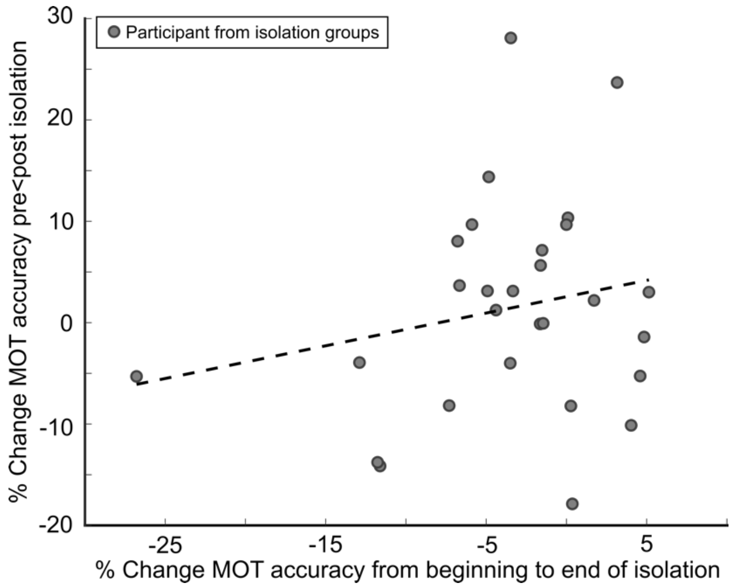


**Supplemental Figure 4:** Percentage change in attended visual field from pre- to post-intervention correlated with percentage change in performance from first to last 10 minutes of training.

This graph demonstrates no relationship between deterioration during training and the performance in the attended visual field for the isolation groups ($r=0.2048$, $p=0.2958$).
